# Stool serology: development of a non-invasive immunological method for the detection of Enterovirus-specific antibodies in Congo gorilla faeces

**DOI:** 10.1101/2020.11.28.402230

**Authors:** Youssouf Sereme, Sandra Madariaga Zarza, Hacène Medkour, Inestin Amona, Florence Fenollar, Jean Akiana, Soraya Mezouar, Nicolas Orain, Joana Vitte, Bernard Davoust, Didier Raoult, Oleg Mediannikov

## Abstract

The incidence of poliovirus has significantly reduced by as much as 99.9% globally. Alongside this, however, vaccine-associated paralytic poliomyelitis has emerged. Recently, a new recombinant virus (Enterovirus C/Poliovirus) was identified in humans working as eco-guards and in gorillas in Democratic Republic of Congo, including one gorilla with polio-like sequelae. A strain of this recombinant virus (Ibou002) was also isolated from gorilla faeces. In order to assess the potential role of poliovirus infection, we have developed and optimised a protocol, based on the lyophilisation and solubilisation of small volumes of stool extracts, to detect specific antibodies. First, total immunoglobulins was detected in the concentrated stool extracts. Specific antibodies were then detected in 4/16 gorilla samples and 2/3 human samples by western blot using both the polio vaccine antigen and the Ibou002 antigen and by ELISA using the polio vaccine antigen. Humoral responses were greater with the Ibou002 antigen. We therefore suggest that this recombinant virus could lead to a polio-like disease in the endangered western lowland gorilla. The development of a non-invasive method to detect microorganism-specific immunoglobulins from faecal samples opens up new perspectives for the exploration of humoral responses of pathogens in animals and a greater understanding of zoonotic infectious diseases.

## INTRODUCTION

Poliovirus is a virus belonging to the Picornaviridae family and the *Enterovirus* genus (Enterovirus C), which causes poliomyelitis in humans.^1^ The poliovirus genome is a 7.5 kilobase single-stranded RNA surrounded by a capsid, composed of four proteins (VP1, VP2, VP3, and VP4).^2^ There are three poliovirus types (1, 2, and 3) and immunity to one type does not protect against the two others.^3^

The global polio eradication programme was initiated in 1988 with the use of a live-attenuated oral poliovirus vaccine. It resulted in the number of wild poliovirus cases in the world declining by more than 99.9%, from an estimated 350,000 cases in more than 125 endemic countries to 175 wild cases in 2019 (https://www.who.int/health-topics/poliomyelitis#tab=tab_1). Wild poliovirus types 2 and 3 were officially certified as having been eradicated in 2015 and 2019, respectively.^4,5^ The non-circulation of wild poliovirus type 1 has been reported in Nigeria since 2016.^14,15^ On 25 August 2020, the WHO African region was certified as being free of wild poliovirus, leaving only two countries in the Eastern Mediterranean Region reporting endemic circulation of wild poliovirus type 1 (https://www.who.int/wer/2020/wer9541/en/).^6,11–13^ In parallel, however, another challenge arose with the emergence of vaccine-associated paralytic poliomyelitis.^6^ In the past 12 months, a total of 132 wild virus cases have been reported in Afghanistan (61) and Pakistan (142) (http://polioeradication.org/polio-today/polio-now/this-week), while 605 cases of vaccine-associated paralytic poliomyelitis have been detected worldwide, including in 19 African countries (406 cases), Afghanistan (101), Pakistan (80), Malaysia (1), the Philippines (1), and Yemen (16). It is notable that of these 605 vaccine-associated cases, 424 originated from non-endemic countries and 181 from known endemic countries.^6–10^ It has also been reported that 5-10% of polio deaths worldwide are now due to cVDPVP^6^.

The transmission route of poliovirus is faecal-oral. Humans remain the natural host for the disease, but it can also infect primates and monkeys.^1,3^ There is a positive correlation between zoonotic diseases and the emergence of infectious diseases in humans.^16^ It has been shown under experimental conditions that hominoids and monkeys can be infected with poliovirus.^17,18^ A 2005 study in Madagascar demonstrated the circulation of vaccine-derived recombinant poliovirus strains, which are recombinant Enterovirus-C viruses, and their ability to lead to a “polio-like” disease in humans.^19^

A study conducted by Harvala *et al.* in Cameroon and the Democratic Republic of Congo (DRC) screened for enteroviruses by PCR in chimpanzee and gorilla faecal samples. It demonstrated both the circulation of genetically divergent variants of enteroviruses in apes and monkeys as well as one strain (EV-A89) that they share with local human populations.^20^ In addition, the *Enterovirus* species described in humans and other species as a potential source of emerging infectious diseases in humans has also been detected in monkeys.^21,22^

Previously, our team reported the presence of a new Enterovirus C which is highly similar to Coxsackievirus as well as similarity with poliovirus 1 and 2, and associated with flaccid paralysis in the chimpanzee virus EV-C99 strains^21^ in the 3C, 3D, and 5’UTR regions.^23^ We hypothesised that a male gorilla living alone in the Lésio-Louna-Léfini Nature Reserve (Republic of the Congo), could be suspected of having a poliovirus infection, based on clinical observations including facial paralysis. Our aim was to develop and to optimise a non-invasive experimental method to assess this hypothesis using gorilla faecal samples to identify specific antiviral antibodies.

## RESULTS

### Differentiation of the faeces of individual gorillas

A previous study by our team had confirmed that each faeces sample corresponded to an individual gorilla.^2.3^ A code was given to each of the 16 gorillas. Twelve had a G0 code followed by a number from 1 to 12, and four had a GNLL code followed by a number from 1 to 4.

### Selection of the stool preparation protocol

Using protocol 1, based on raw stool extracts, very low concentrations of immunoglobulins were detected (Supplementary Figure 1). Using protocol 2, based on purification with protein G and peptin M, we noticed a loss of immunoglobulin concentrations after purification (data not shown). Protocol 3, based on the lyophilisation of the filtered stool extract followed by its reconstitution in a small volume, led to the best results, as shown in Figure 1. In the gorilla stools, the median concentration was 223.5 ng/ml with an interquartile range (IQR) of 172.8-272 observed for IgA, and 135 ng/ml with an IQR of 80.23–171.7 for IgG. In the human faeces, the median concentration was 1612.1 ng/ml with an IQR of 812.12–1965.30 observed for IgA, and 639.3 ng/ml with an IQR of 503.3–698 for IgG. The median IQR was the 25–75 percentile (Figure 2). The latter was therefore selected to continue analyses on the stool.

**Figure 1.**
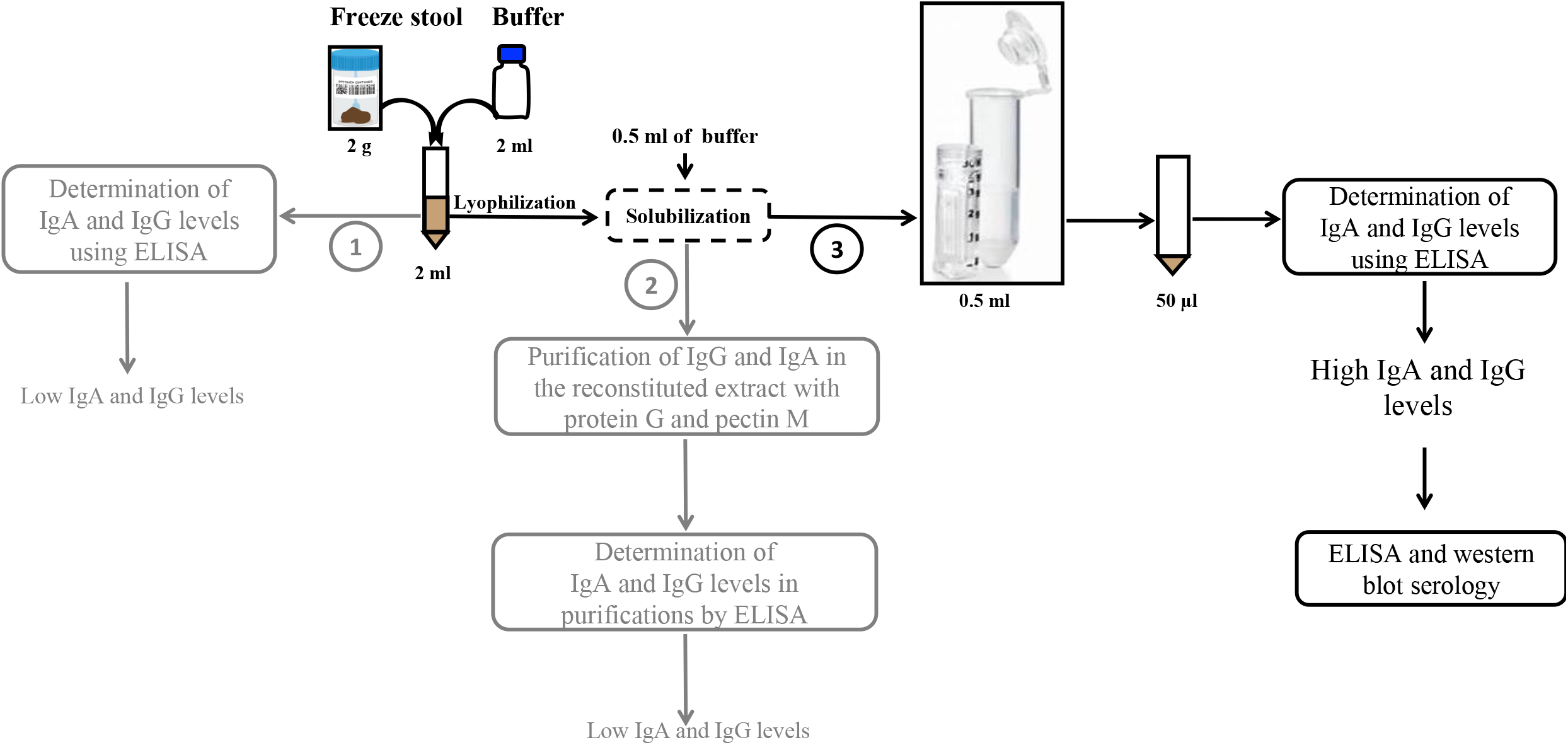
Schematic representation of the three protocols evaluated for protein extraction from stools up to evaluation of immunoglobulins by ELISA and western blot. Protocol 3, indicated in black, is the one that was retained.

**Figure 2.**
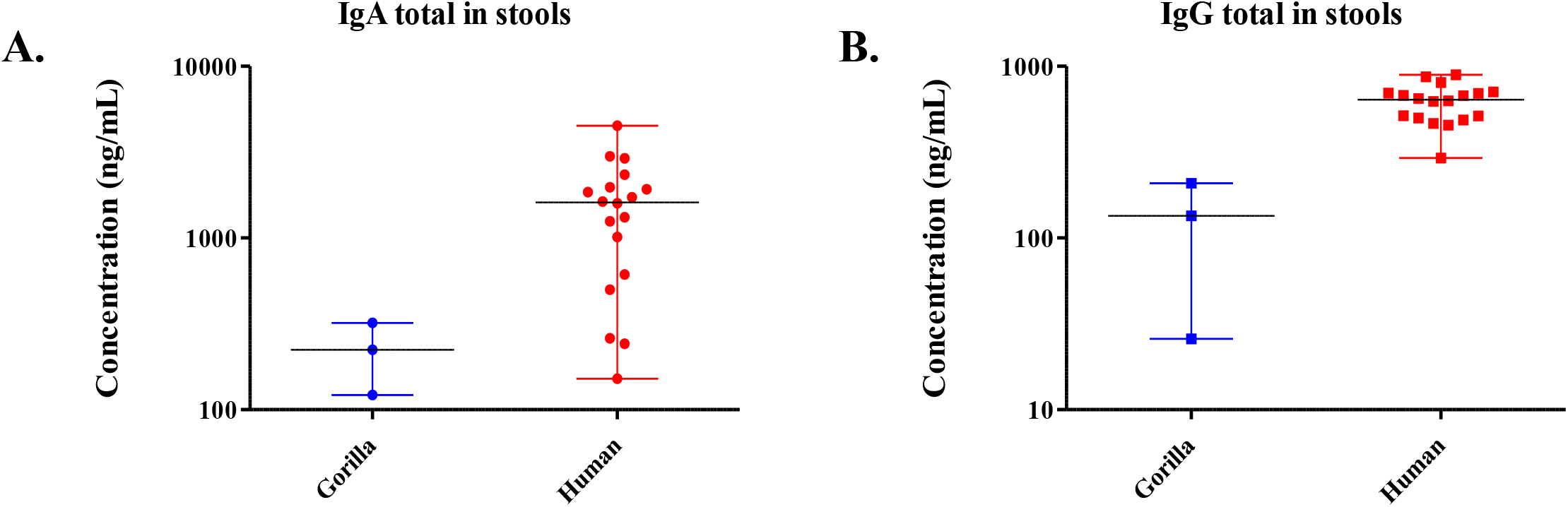
Evaluation of total IgA (A) and IgG (B) concentrations (ng/mL) in stools of three gorillas and nineteen humans by ELISA. Concentrations (ng/mL) gorilla and human total IgA (A) and IgG (B) are expressed as median and interquartile ranges (IQR). The stool concentration factor and median and IQR were not included in our calculations, and the median and IQR were calculated by restricting the results above the lower LOQ (limit of quantification) for each analyte.

### Quantification and detection of poliovirus-specific immunoglobulins by ELISA and western blot

#### Quantification of poliovirus-specific immunoglobulins (IgA-IgG) by ELISA

The poliovirus-IgG/IgA evaluation of 16 gorilla and 3 human stool samples was carried out by ELISA. The IgG/IgA level was represented by the OD at 450 nm. A sample was concluded as positive for IgG / IgA when the OD value was greater than twice the negative control value for the kit used (OD positive > 0.35). Specific antibodies were detected in four of 16 (25%) gorilla samples (G01, G02, G03, and G07), and in two of the three human faecal samples (Ibou2 and Ibou3) (Figure 3). In addition, the recombinant Enterovirus C (strain Ibou002) was isolated from one of the two eco-guards (Ibou3) who were positive for poliovirus.^22^

**Figure 3.**
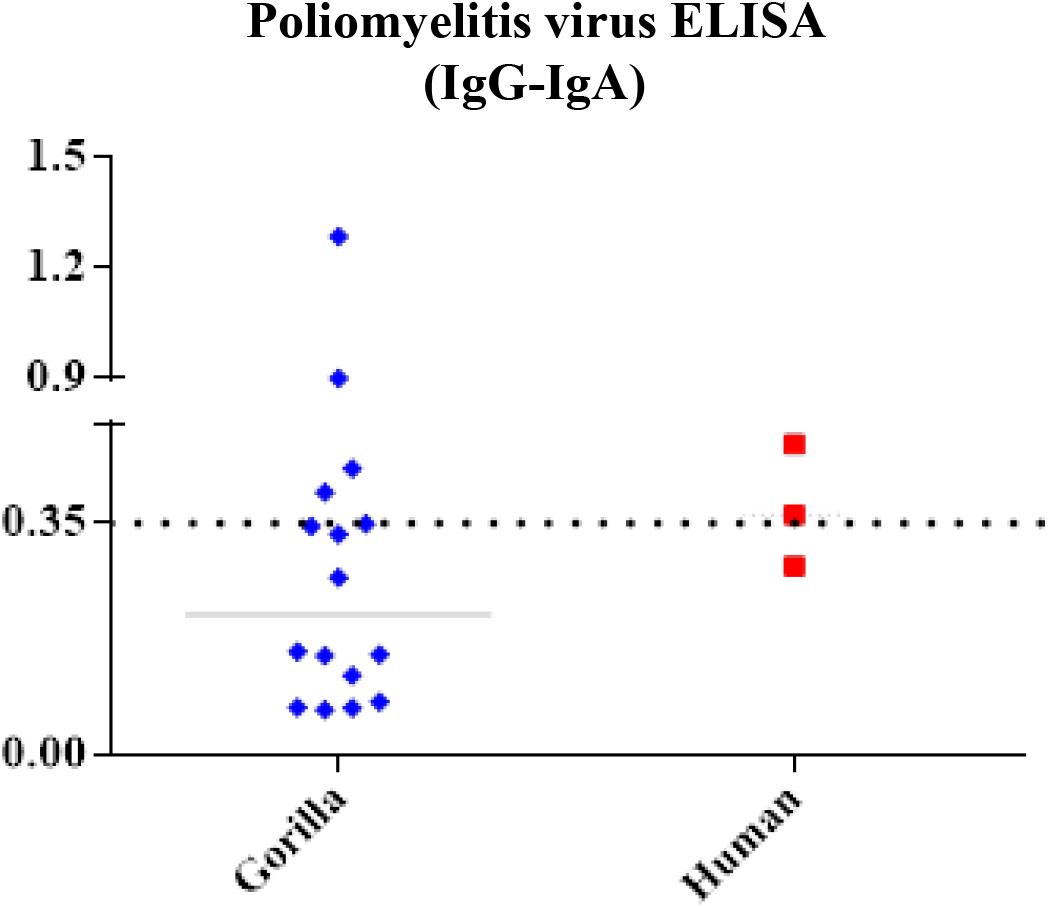
Evaluation of poliovirus IgG/IgA by ELISA. Graph representing the specific IgG/IgA (OD) level of all samples, measured at a wavelength of 450 nm. A sample was concluded as being positive for IgG/IgA when the OD value was greater than twice the negative control value for the kit used (OD positive > 0.35).

#### Detection of poliovirus- and enterovirus C-specific immunoglobulins by western blot Protein profiles of both antigens (Imovax vaccine and strain Ibou002)

The protein profile was performed for both antigens (Imovax vaccine and strain Ibou002). Different protein profiles were identified for each antigen by SDS-PAGE. The vaccine displayed five bands possibly indicating five proteins with molecular weights ranging from 20-35 kDa. We found seven bands in the antigen purified from the virus, which correspond to a molecular weight of between 28 and 130 kDa (Figure 4).

**Figure 4.**
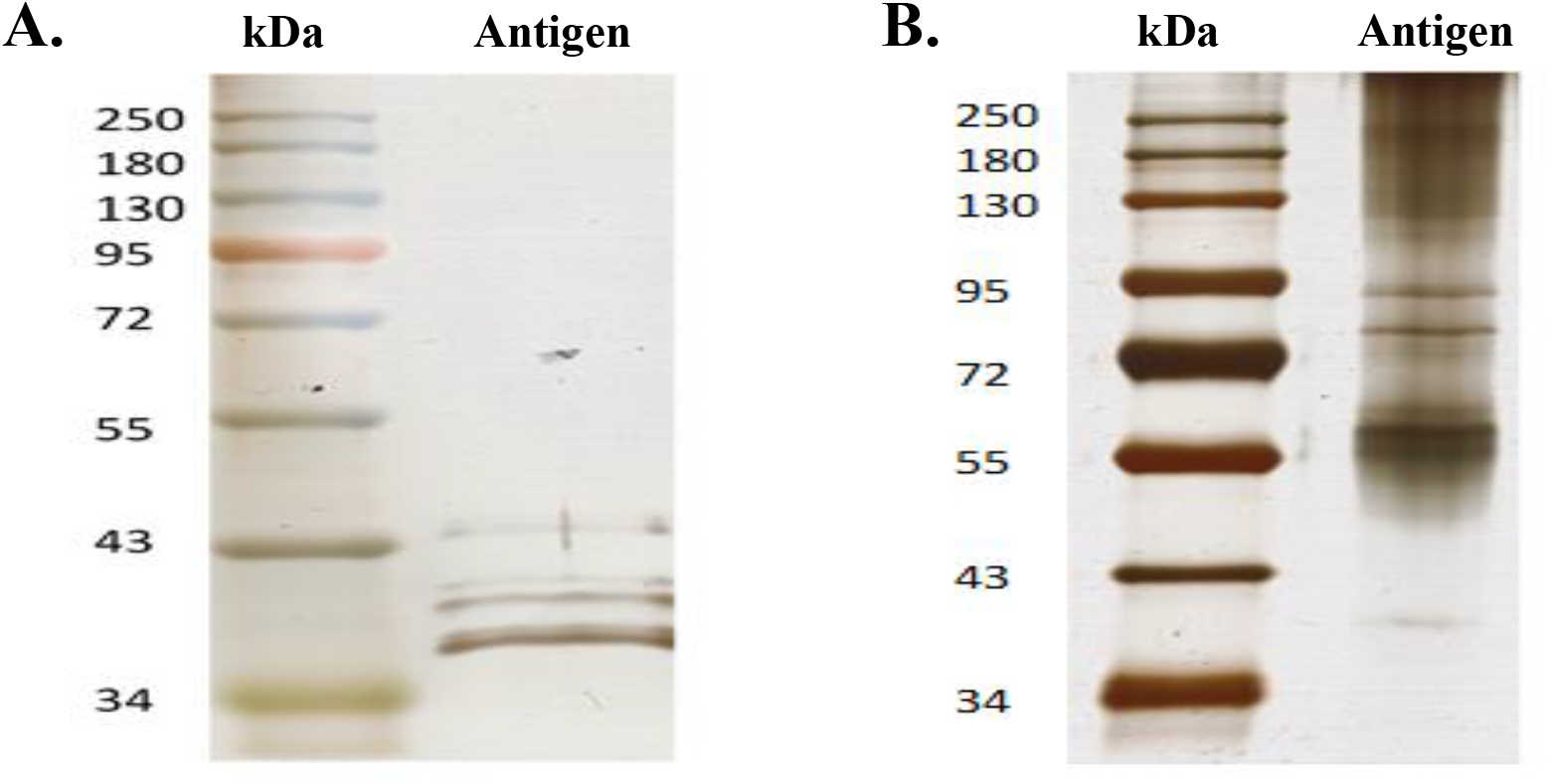
Silver stain SDS-PAGE analyses for poliovirus vaccine (A) and antigen of Ibou002 virus strain (B) proteins. In both profiles, the left lane corresponds to the standard molecular weight and the right lane to the antigen; 50 μg of poliovirus vaccine and 30 μg of antigen of Ibou004 virus strain were run in a 12% polyacrylamide gel.

#### Western blot

The immunoglobulin evaluation by western blot showed positive samples for both antigens source (poliovirus vaccine and Ibou002 virus strain), but with different band profiles between them. Bands were detected in four samples (G01, G02, G03, and G07) from 16 samples of gorilla faeces, and two samples (Ibou2 and Ibou3) from three human faeces, which were previously detected as positive by ELISA (Figure 5). We also noted that all samples with bands from the vaccine antigen also showed bands for the virus isolated, but at different molecular weight depending on the antigen source (14 kDa for vaccine antigen, and 60 kDa and 116 kDa for antigen Ibou002 virus strain).

**Figure 5.**
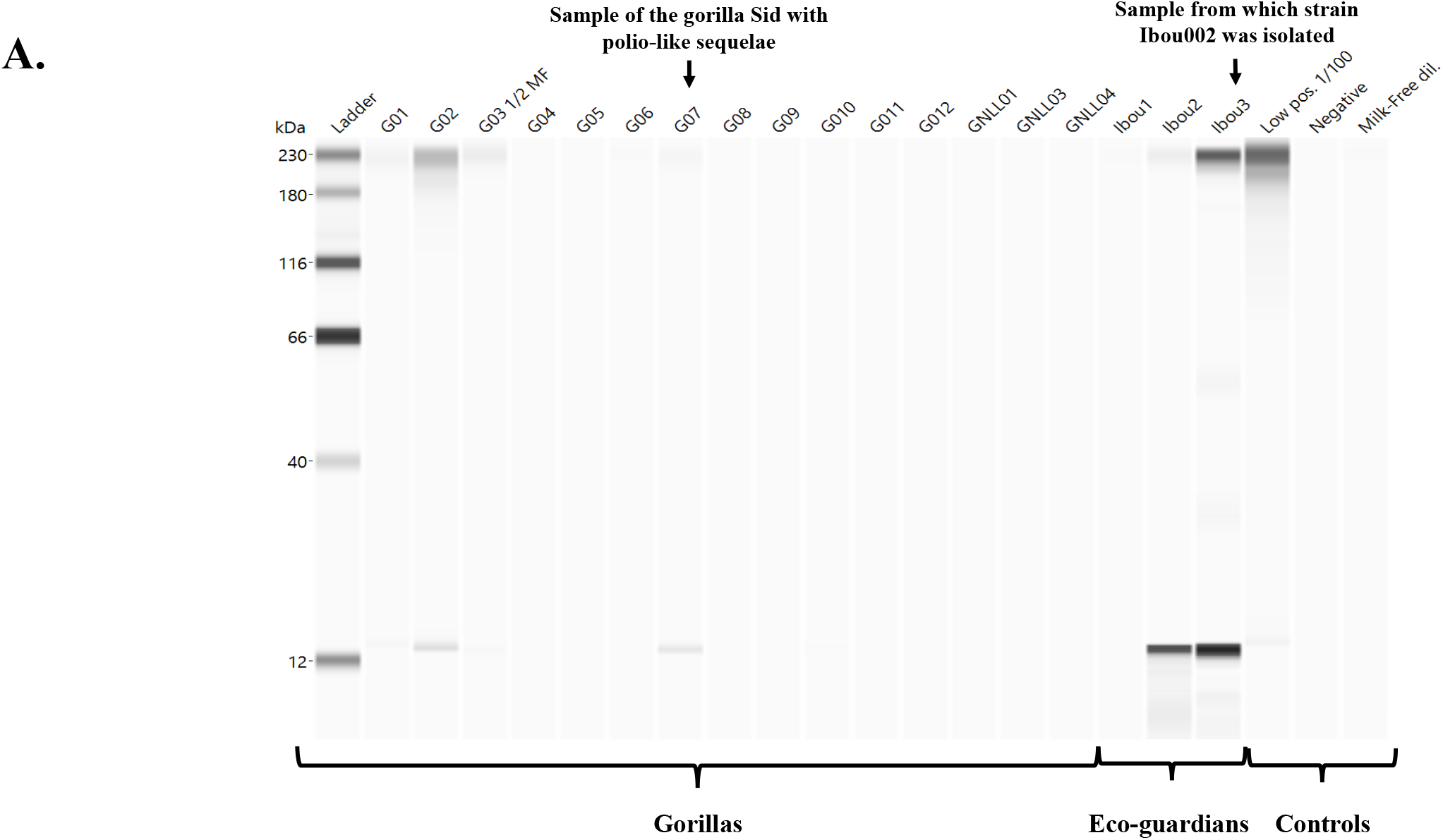
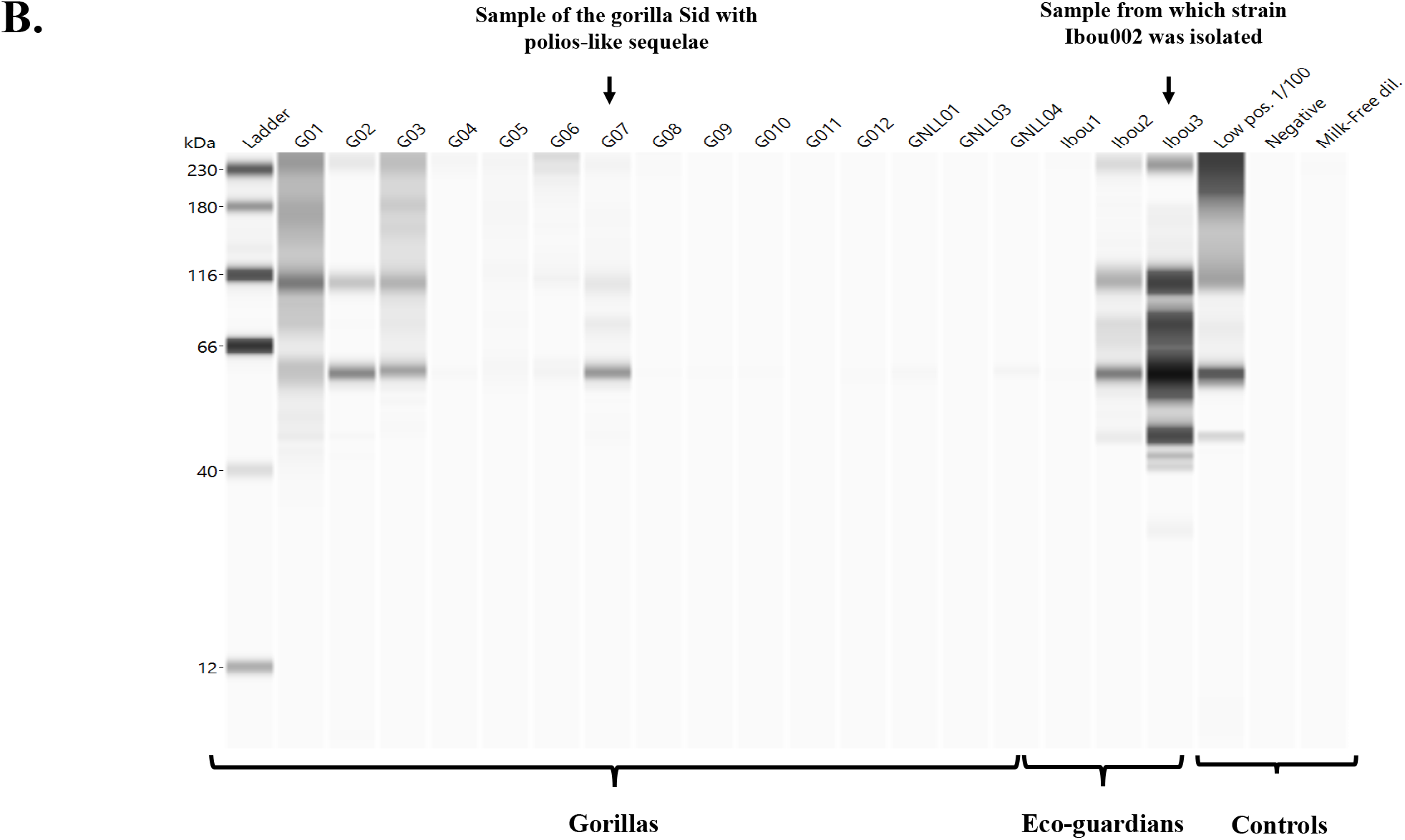
Western blot Jess^TM^ using two different antigens, poliovirus vaccine (A) and Ibou002 virus strain (B). Anti-human IgG/IgM/IgA antibody detection was performed for 16 gorilla stools, three human stools, and one control positive serum (non-diluted and diluted 1:100).

## DISCUSSION

In this study, we developed a non-invasive method for screening animal immune responses using fresh stool samples. We also provided a proof of concept on the feasibility of exploring immunity from faeces. The detection of the total antibody levels (IgA / IgG) in concentrated stool extracts via the developed protocol confirms the possibility of using these samples for the detection, quantification and characterisation of specific antibodies by ELISA and western blot. The very high concentrations of IgA in faeces can be explained by the microbial colonisation of the gut that triggers the production of IgA by the gut associated lymphoid tissue (GALT).^25^ This supports our decision to use an anti-IgA, IgG mixture in our ELISA and western blot techniques to detect the maximum amount of specific antibodies. In addition, we did not look for specific IgM because the concentration of total IgM in the digestive mucosa is very low or even zero.^26^

By testing a working hypothesis that circulating (probably vaccine-associated) poliovirus^19^ caused poliomyelitis-like disease in gorillas, we attempted to identify specific antipoliovirus antibodies in the stools of 16 gorillas and three eco-guards working in the reserve. Using our original method of antibody extraction, we detected antibodies binding to the poliovirus antigen in faecal samples by ELISA^27^, enabling us to identify positive samples in four of the 16 gorillas and two of the three eco-guards. Such results in humans were not surprising, as the majority of the population has been vaccinated against poliomyelitis in this area. However, the positive results in four gorillas, including Sid, the gorilla with sequelae, were more surprising, indicating a potential circulation of the poliovirus or a closely-related Enterovirus in this community. We then performed a western blot by Jess^TM^ to characterise the specific antibodies. Western blot analysis of antibodies from gorilla faeces detected a single band of around 14 kDa using the antigens extracted from poliovirus vaccine. These data do not allow us to confirm the gorilla’s immunisation from poliovirus due to the absence of antibodies against the main proteins of the poliovirus, namely VP1, VP2, and VP3. In addition, PCR testing for poliovirus RNA was negative.^22^

Based on the circulation of recombinant Enterovirus C^22^ having at least three genes very close to the poliovirus in this geographical area, we then hypothesised that Sid the gorilla may have been suffering from the sequelae of a polio-like illness caused by this virus. Indeed, these polio-like diseases can be caused in humans by chimeric/recombinant viruses.^19–21^ These diseases often produce typical sequelae in the form of flaccid paralysis and muscle atrophy. In order to verify this hypothesis, we prepared the antigen from the recombinant Enterovirus C (Ibou002 strain) previously isolated from a stool sample from an eco-guard in the Lésio-Louna Reserve. We noticed that the four samples of gorilla faeces that showed bands with the vaccine antigen were the same samples that showed bands with the antigen from the recombinant Enterovirus C. This concordance was also observed with the two human faeces samples. Western blot analysis detected specific immunoglobulins to different viral proteins. Indeed, two 60 kDa and 110 kDa molecular weight bands were present in all positive samples from gorillas and humans with the Ibou002 strain antigen, and one 14 kDa band was also present in all positive samples with the vaccine antigen. Evaluation of the western blot results showed that the positive specimens for the two antigen sources (polio vaccine and Ibou002 virus strain) were the same but with different profiles. Thus, we hypothesise that the recombinant Enterovirus C that we recently isolated from an eco-guard may be in wide circulation and was probably responsible for the immunisation of the humans and the gorillas.

Given the presence of typical post-polio sequelae in Sid the gorilla, the presence of specific antibodies binding to the Ibou002 virus strain and the recombinant nature of this virus, we suggest that this new virus could cause a polio-like disease in the endangered western lowland gorilla. We also believe that the detection of a band in western blot with the poliovirus antigen is the result of a non-specific cross-reaction between the antibodies induced by the recombinant Enterovirus C strain Ibou002 and the genetically related poliovirus. Another interesting finding is the detection of specific anti-recombinant Enterovirus C Ibou002 antibodies in the same stool sample from the person from whom the virus was isolated. This may indicate a long persistence of the virus in the human organism and the absence of protective immunity.

## CONCLUSION

As a result of the non-invasive immunological method developed using faeces, we were able to assess the presence of specific antibodies against a recombinant Enterovirus C in gorillas and eco-guard in the same reserve. This method, which does not require medical or invasive procedures for the collection of samples and which allows for the detection of specific antibodies, opens numerous prospects for application in zoonotic infectious diseases and could revolutionise the screening of animals for important emerging infections, such as Ebola fever, rabies, and coronavirus infections.

## MATERIALS AND METHODS

### Collection of gorilla faeces samples

Fresh faecal samples were collected from western lowland gorillas *(Gorilla gorilla gorilla)* living in semi-captivity and in the wild in the Lésio-Louna-Léfini Gorilla Nature Reserve, located in the northern Pool Department in the Republic of Congo. A total of 16 gorilla faecal samples were collected. Samples were collected from Iboubikro (Lat: 03.27040; Long 15.47071) from semi-captive young gorillas and from Abio2 (Lat 03.094; Long 15.52633) from wild gorillas. Samples were collected in individual pots (Labelians, Nemours, France) and 50 ml Falcon tubes (Dutscher, Brumath, France). At each collection time and for each sample, information relating to the GPS position of the location, the estimated decomposition time of the faeces as well as the morphological and/or physical aspect of the sample was recorded. During the sample collection, we observed a male gorilla named Sid living alone on an island in front of the Abio station in the Lésio-Louna-Léfini Nature Reserve, where he had been placed after being sick. We observed clear signs of facial myodystrophy and paresis and upper limbs paresis (Figure 6). Three faecal samples were collected from three people working on the site as “eco-guards”. Each sample was given an anonymous identification code (Ibou1, Ibou2, and Ibou3). A poliovirus serology-positive control serum (Poliomyelitis virus kit, GenWay, San Diego, California, USA) was also used as a control. Research and administrative authorisations for the collection of samples were granted by the Congolese Ministry of Scientific Research and Technological Innovation (N°003/MRSIT/DGRST/DMAST), the Ministry of Forestry Economy and Sustainable Development, represented by the Congolese Agency for Wildlife and Protected Areas (N°0134/ACFAP-DTS) and (N°94/MEFDD/CAB/DGACFAP-DTS), and the Ministry of Health and Population (N°000220/MSP-CAB-17 and N°208/MSP/CAB.15).

**Figure 6.**
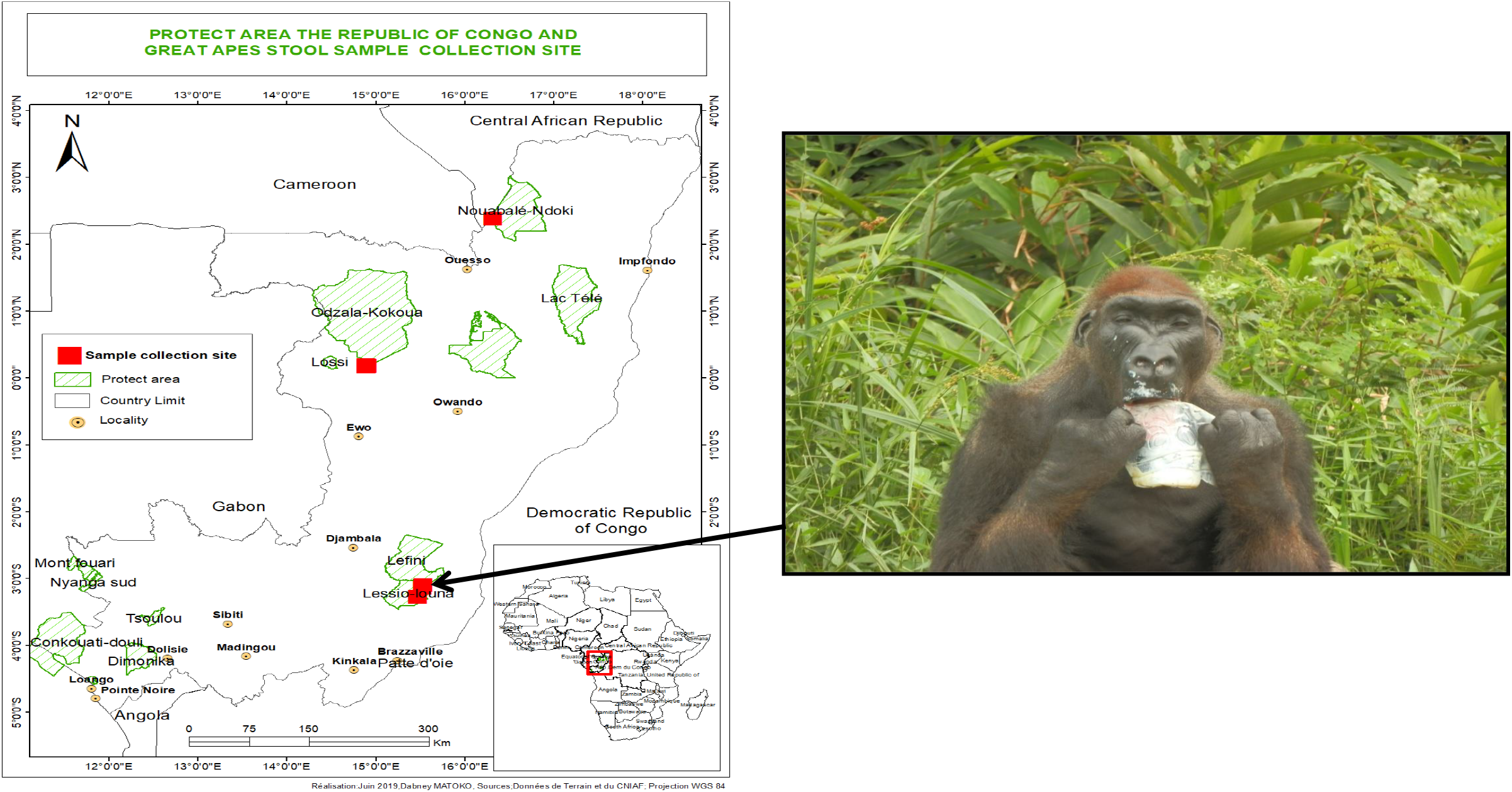
Geographic map of the collection site and the gorilla named “Sid” who presented myodystrophy and paresis on the face. He lives alone on an island in front of the Abio station in the Lesio-Louna nature reserve (Republic of Congo).

### Concept and optimisation of the methodological approach

The principle of our approach initially consisted in developing a method of extracting immunoglobulins which makes it possible to obtain structural and functional proteins for humoral evaluation. The extraction buffer contained a protease inhibitor cocktail to inhibit the general degradation of proteins from stool extracts, tissues or cells by endogenous or exogenous microorganism proteases and to study processes that involve blocking the activity of specific proteases. Three gorilla and 19 human stool samples were used to optimise the concept. Different stool treatment protocols were tested for the detection of immunoglobulins (IgG and IgA), mainly: (1) filtration of stool extracts from 2 g of stools plus 2 ml of buffer; (2) lyophilisation of the filtered extract, reconstituted in 500 μl of buffer and then purification of IgG and IgA in the reconstituted extract with protein G and pectin M; and (3) lyophilisation of the filtered extract, reconstituted in a small volume of buffer (500 μl) then concentrated by the Amicon^®^ Ultra-1. After evaluation of the data, a final protocol was adopted (Figure 1).

### Experimental protocol adopted for stool protein extraction

Two grams of each sample were solubilised in 2 μl of an extraction buffer consisting of phosphate buffered saline solution supplemented with 4-(2-aminoethyl)-benzenesulfonyl fluoride 4 mM, bestatin 0.26 mM, E-64 28 μM, leupeptin 2 μM and aprotinin 0.6 μM, pH 7.4, and SIGMAFAST™ Protease Inhibitor Tablets (Sigma-Aldrich, Saint-Louis, Missouri, USA). The stool-buffer mixture was incubated for 15 minutes at room temperature, vortexed for 1 minute, and then centrifuged at 1,500 rpm for 15 minutes at 4°C. The supernatant was lyophilised for 24 hours and then re-solubilised in 1 ml of distilled water. Then, 500 μl of resolubilised stool was concentrated using Amicon^®^ Ultra-15 3k centrifugal filters (Merck, Saint-Romain, France) for further studies (Figure 1). Concentrations of total immunoglobulins (IgA and IgG) were evaluated by ELISA (Abcam, Paris, France) to validate the method of protein extraction from stools (Figure 1).

### Quantification of anti-poliovirus immunoglobulins by ELISA

Specific immunoglobulins were identified (IgG and IgA) by ELISA according the manufacturer’s instructions (Poliomyelitis virus kit, GenWay, San Diego, California, USA). The polio antigen derived from the human pathogenic poliovirus types, specifically type 1 (Brunhilde), type 2 (Lansing), and type 3 (Leon) was bound in the wells of the plate. The diluted samples or standards were then deposited. After one hour of incubation at room temperature, the plate was rinsed with wash buffer. The anti-human IgG peroxidase conjugate was then added and incubated for 30 minutes, followed by a wash step. The substrate solution was then added and incubated for 20 minutes. The reaction was stopped by adding a stop solution. The resulting dye was measured using a Thermo Fisher spectrophotometer (Uppsala, Sweden) at a wavelength of 450 nm.

### Characterisation of enterovirus-specific antibodies by western blot from faecal samples

#### Preparation of antigen from the polio vaccine

The Imovax Polio vaccine (Sanofi, Antony, France) was used as an antigen source for the characterisation of poliovirus-specific immunoglobulins by western blot. The vaccine is a sterile suspension of poliovirus types 1 (Mahoney), 2 (MEF1), and 3 (Saukett), which are grown on Vero cells, purified, and then inactivated with formaldehyde. Four doses of Imovax Polio were treated with the TS lysis buffer and fractionated by sonication to release the antigen molecules. The antigen suspension was lyophilised and re-solubilised in 100 μl of distilled water (Figure 7). The concentrated antigen was measured by Bradford protein assay (Bio-Rad, Hemel Hempstead, UK), and an SDS-PAGE was performed with 50 μg of the purified antigen and 12% of polyacrylamide.

**Figure 7.**
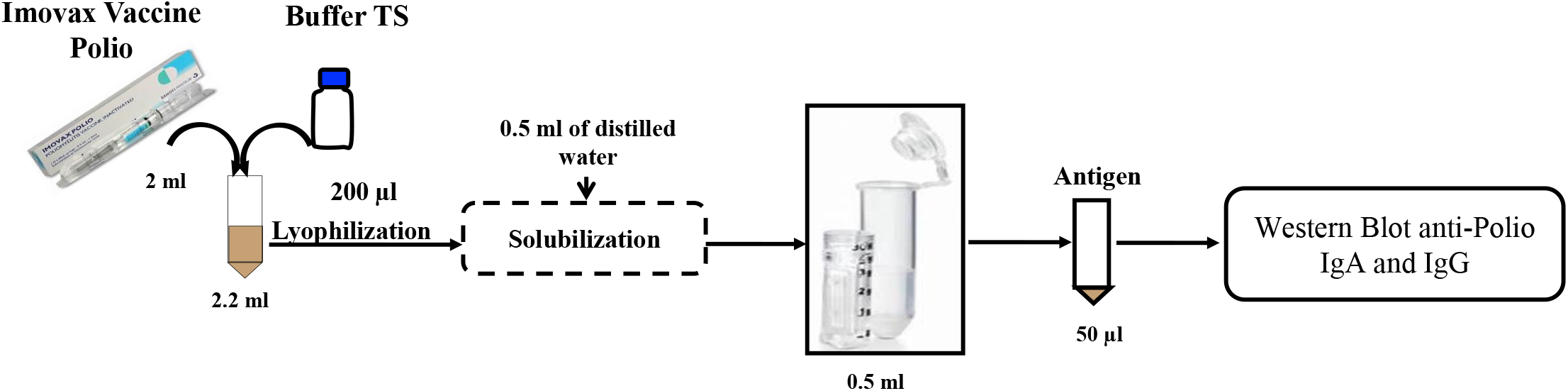
Protocol for the extraction of antigens from the polio vaccine (Imovax^®^) for analysis by western blot.

#### Preparation of antigen from enterovirus strain Ibou002

The strain Ibou002^23^ of recombinant enterovirus C, originally isolated from an eco-guardian in the Lésio-Louna Nature Reserve, was replicated for seven days in MRC5 cells line (ATCC^®^ CCL-171™). The virus-infected MRC5 cells were collected and centrifuged at 700 × g for 15 minutes (Eppendorf Centrifuge 5810R). Supernatant was filtered by serial filtration steps to eliminate cells and cellular debris, passing through 0.8 μm, 0.45 μm, and 0.2 μm filters. Virus particles were then precipitated in 10% PEG-8000 and 2.2% NaCl at 4°C overnight, in agitation. The precipitates were centrifuged at 10,000 × g (Sorvall Evolution, SLA-3000 Recent-1 rotor, Thermo Fisher) for 30 minutes at 4°C, and the virus pellet was then resuspended in HEPES saline solution (9 g NaCl, 10 mL HEPES 1M, 990 mL distilled H2O). The resuspended virus was purified using a 30% (wt/vol) sucrose cushion at 100,000 × g (Sorvall Discovery 90SE, Thermo Fisher) for two- and-a-half hours at 4°C. The virus pellet was washed in HBSS (Thermo Fisher), and centrifuged at 100,000 × g (Beckman SW 32 Ti rotor) for 30 minutes at 4°C. Finally, the viral particles were resuspended in HEPES saline and then fractionated with TS buffer (7 M Urea, 2 M Thiourea, 4% CHAPS) to release the antigen. The released antigen was then concentrated with the Amicon 3 kDa filter (Merck KGaA, Darmstadt, Germany) before being analysed by SDS gel electrophoresis, and western blot. The concentration of the antigen was measured, and 30 μg of the Purified product was loaded onto 12% SDS-polyacrylamide gel for analysis.

#### Western blot

Western blot was performed using Jess^TM^ Simple Western system (Protein Simple, San Jose, California, USA) according to the manufacturer’s protocol.^24^ This technique is an automated capillary size separation and nano-immunoassay system for performing a gel-free, blot-free, and hands-free capillary-based immunoassay that integrates and automates the entire protein separation and detection process with home-made antigens. To quantify the absolute response to viral proteins from samples, we followed the manufacturer’s standard method for 12-230 kDa Jess^TM^ separation module (SM-W004). First, the poliovirus antigen (0.24 μg/μL) was mixed with fluorescent 5X master mix (Protein Simple) to achieve a final concentration of 0.20 μg/μL in the presence of fluorescent molecular weight markers and 400 mM dithiothreitol (DTT, Protein Simple). This preparation was then denatured at 95°C for five minutes. The ladder (12-230 kDa) and viral proteins were separated in capillaries as they migrate through a separation matrix at 375 volts. A protein simple proprietary photoactivated capture chemistry was used to immobilise separated viral proteins on the capillaries. Non-diluted faecal samples were added and incubated for 30 minutes. After a wash step, goat HRP-conjugated anti-human IgG/IgM/IgA antibodies (Jackson ImmunoResearch, Ely, UK) diluted at 1:250 were added for 30 minutes. Finally, the chemiluminescent revelations of the ladder and samples were established with peroxide/luminol-S (Protein Simple). The digital image was analysed with Compass for SW software (v 4.1.0, Protein Simple), and quantified data of the detected proteins was reported, including the molecular weight, chemiluminescence intensity, and signal/noise ratio. Western blot analyses were performed according to this protocol, using (1) the polio vaccine and (2) the purified Ibou002 virus strain as antigens.

## ACKNOWLEDGEMENTS

We thank Lucile Pinault and Rita Jaafar for their help and advice in the development of the experiments. This work was supported by the French Government under the “Investissements d’avenir” (Investments for the future) programme managed by the “Agence Nationale de la Recherche” (reference: 10-IAHU-03). Research and administrative authorisations for the collection of samples were granted by the Congolese Ministry of Scientific Research and Technological Innovation (n°003/MRSIT/DGRST/DMAST), the Ministry of Forestry Economy and Sustainable Development, represented by the Congolese Agency for Wildlife and Protected Areas (N° 0134/ACFAP-DTS) and (N° 94/MEFDD/CAB/DGACFAP-DTS), and the Ministry of Health and Population (N°000220/MSP-CAB-17 and N°208/MSP/CAB.15). YS, SZM, HM are supported by “Fondation Méditerranée Infection” doctoral positions. SM was supported by a post-doctoral “Fondation Méditerranée Infection” fellowship.

## AUTHOR CONTRIBUTIONS

YS conducted the experiments, analysed the data and wrote the paper; SMZ, HM, JV, NO and SM helped conduct the experiments and participated in the writing of the paper; JA, BD and IA contributed to the collection of samples; OM, DR, FF designed the project, participated in the writing of the paper and provided great support carrying out the experiments.

---

**Supplementary figure 1.** Quantification of immunoglobulins (IgG and IgA) in gorilla faeces with three different faecal treatment protocols (1) Filtration of stool extract from 2 g of stools plus 2 ml of buffer; (2) Lyophilisation of the filtered extract, reconstituted in 500 μl of buffer and then purification of IgG and IgA in the reconstituted extract with protein G and pectin M; (3) Lyophilisation of the filtered extract, reconstituted in a small volume of buffer (500 μl) then concentrated by the Amicon^®^ Ultra-1.

## References

1. Fields virology. (Wolters Kluwer Health/Lippincott Williams & Wilkins, Philadelphia, 2007).

2. Blondel, B. et al. Genetic evolution of poliovirus: success and difficulties in the eradication of paralytic poliomyelitis. Med. Trop. (Mars) 68, 189–202 (2008).

3. Pelletier, J. & Sonenberg, N. Internal initiation of translation of eukaryotic mRNA directed by a sequence derived from poliovirus RNA. Nature 334, 320–325 (1988).

4. Khan, F. et al. Progress Toward Polio Eradication – Worldwide, January 2016-March 2018. Morb. Mortal. Wkly. Rep. 67, 524–528 (2018).

5. Weekly epidemiological record: Progress towards poliomyelitis eradication. World Health Organization (WHO) (2019).

6. Javelle, E. & Raoult, D. Antibiotics against poliovirus carriage: an additional tool in the polio endgame? Clin. Microbiol. Infect. 26, 542–544 (2020).

7. Mbaeyi, C. et al. Update on Vaccine-Derived Poliovirus Outbreaks — Democratic Republic of the Congo and Horn of Africa, 2017-2018. Morb. Mortal. Wkly. Rep. 68, 225–230 (2019).

8. Saleem, A. F. et al. Immunogenicity of Different Routine Poliovirus Vaccination Schedules: A Randomized, Controlled Trial in Karachi, Pakistan. J. Infect. Dis. 217, 443–450 (2018).

9. Chandler, J. Fighting a polio outbreak in Papua New Guinea. The Lancet 392, 2155–2156 (2018).

10. Roberts, L. Polio eradication campaign loses ground. Science 365, 106–107 (2019).

11. Akhtar, R. et al. Genetic Epidemiology Reveals 3 Chronic Reservoir Areas With Recurrent Population Mobility Challenging Poliovirus Eradication in Pakistan. Clin. Infect. Dis 71, e58–e67 (2020).

12. Kalkowska, D. A., Pallansch, M. A. & Thompson, K. M. Updated modelling of the prevalence of immunodeficiency-associated long-term vaccine-derived poliovirus (iVDPV) excreters. Epidemiol. Infect. 147, e295 (2019).

13. Martinez, M. et al. Progress Toward Poliomyelitis Eradication — Afghanistan, January 2019-July 2020. Morb. Mortal. Wkly. Rep. 69, 1464–1468 (2020).

14. Duintjer Tebbens, R. J. & Thompson, K. M. Polio endgame risks and the possibility of restarting the use of oral poliovirus vaccine. Expert Rev. Vaccines 17, 739–751 (2018).

15. Thompson, K. M. & Duintjer Tebbens, R. J. Lessons From the Polio Endgame: Overcoming the Failure to Vaccinate and the Role of Subpopulations in Maintaining Transmission. J. Infect. Dis. 216, S176–S182 (2017).

16. McArthur, D. B. Emerging Infectious Diseases. Nurs. Clin. North Am. 54, 297–311 (2019).

17. Hsiung, G. D. Diagnostic virology: from animals to automation. Yale J. Biol. Med. 57, 727–733 (1984).

18. Ida-Hosonuma, M. et al. Host range of poliovirus is restricted to simians because of a rapid sequence change of the poliovirus receptor gene during evolution. Arch. Virol. 148, 29–44 (2003).

19. Rakoto Andrianarivelo, M. et al. Reemergence of Recombinant Vaccine Derived Poliovirus Outbreak in Madagascar. J. Infect. Dis. 197, 1427–1435 (2008).

20. Harvala, H. et al. Co-circulation of enteroviruses between apes and humans. J. Gen. Virol. 95, 403–407 (2014).

21. Mombo, I. M. et al. First Detection of an Enterovirus C99 in a Captive Chimpanzee with Acute Flaccid Paralysis, from the Tchimpounga Chimpanzee Rehabilitation Center, Republic of Congo. PLoS ONE 10, e0136700 (2015).

22. Mombo, I. M. et al. African Non-Human Primates Host Diverse Enteroviruses. PLoS ONE 12, e0169067 (2017).

23. Amona, I. et al. Isolation and Molecular Characterization of Enteroviruses from Humans and Great Apes in the Republic of Congo: Recombination within Enterovirus C Serotypes. Microorganisms 8, 1779 (2020).

24. Wu, J., Haitjema, C. H., Heger, C. D. & Boge, A. A proof-of-concept analysis of carbohydrate-deficient transferrin by imaged capillary isoelectric focusing and in-capillary immunodetection. BioTechniques 68, 85–90 (2020).

25. Suzuki, K., Ha, S., Tsuji, M. & Fagarasan, S. Intestinal IgA synthesis: A primitive form of adaptive immunity that regulates microbial communities in the gut. Seminars in Immunology 19, 127–135 (2007).

26. Fadlallah, J. et al. Microbial ecology perturbation in human IgA deficiency. Sci. Transl. Med. 10, eaan1217 (2018).

27. Kummitha, C. M. et al. A sandwich ELISA for the detection of Wnt5a. J. Immunol. Methods 352, 38–44 (2010).

