## Supplemental Data 1 for "Stool serology: development of a non-invasive immunological method for the detection of Enterovirus-specific antibodies in Congo gorilla faeces"

**Supplementary figure 1.** Quantification of immunoglobulins (IgG and IgA) in gorilla faeces with three different faecal treatment protocols (1) Filtration of stool extract from 2 g of stools plus 2 ml of buffer; (2) Lyophilisation of the filtered extract, reconstituted in 500 µl of buffer and then purification of IgG and IgA in the reconstituted extract with protein G and pectin M; (3) Lyophilisation of the filtered extract, reconstituted in a small volume of buffer (500 µl) then concentrated by the Amicon® Ultra-1.

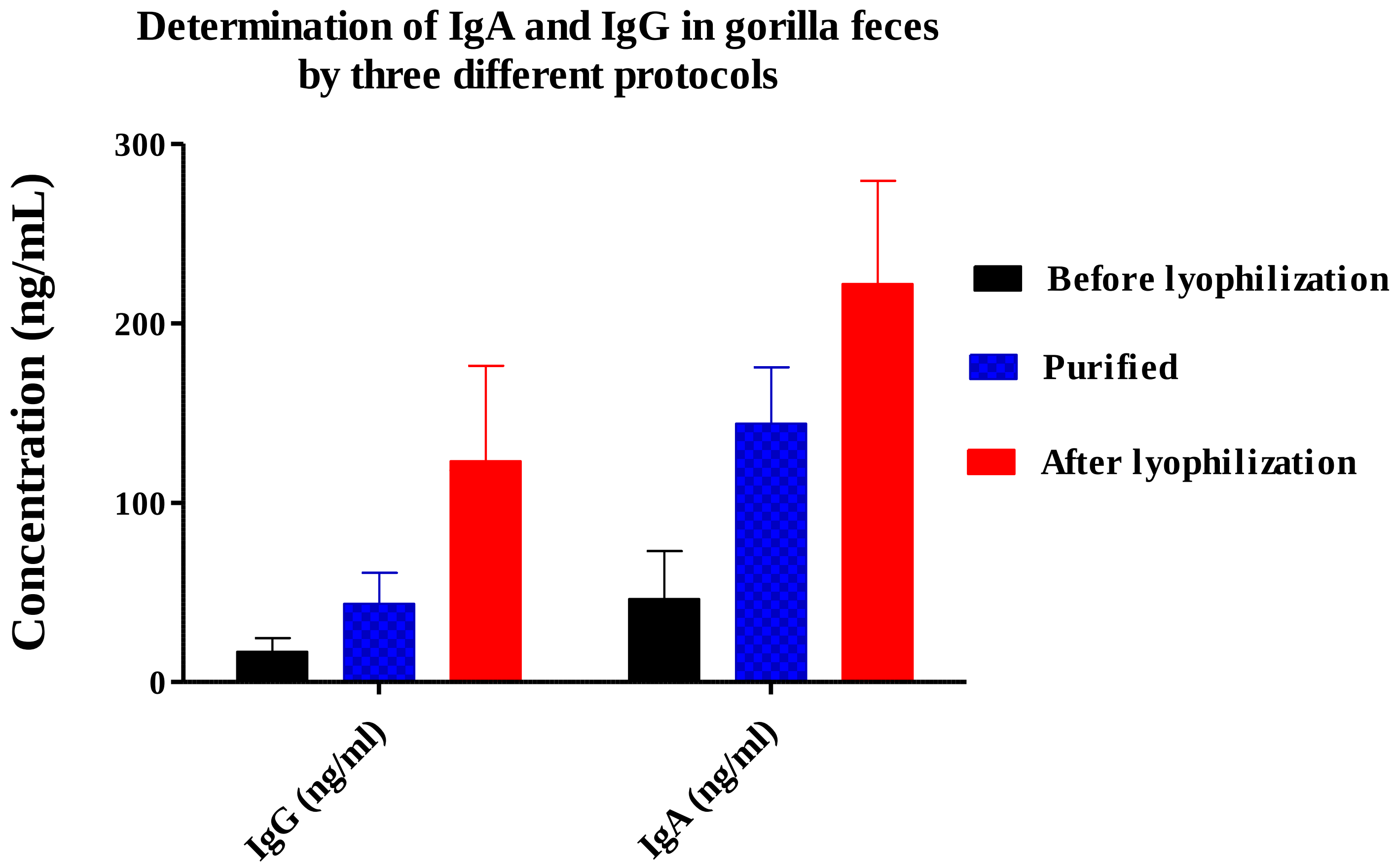
